# Longitudinal Associations Between MicroRNAs and Weight in the Diabetes Prevention Program

**DOI:** 10.1101/2024.06.05.597590

**Authors:** Elena Flowers, Benjamin Stroebel, Xingyue Gong, Kimberly Lewis, Bradley E. Aouizerat, Meghana Gadgil, Alka M. Kanaya, Li Zhang

## Abstract

**OBJECTIVE:** Circulating microRNAs show cross-sectional associations with overweight and obesity. Few studies provided data to differentiate between a snapshot perspective on these associations versus how microRNAs characterize prodromal risk from disease pathology and complications. This study assessed longitudinal relationships between circulating microRNAs and weight at multiple time-points in the Diabetes Prevention Program trial.

**RESEARCH DESIGN AND METHODS:** A subset of participants (n=150) from the Diabetes Prevention Program were included. MicroRNAs were measured from banked plasma using a Fireplex Assay. We used generalized linear mixed models to evaluate relationships between microRNAs and changes in weight at baseline, year-1, and year-2. Logistic regression was used to evaluate whether microRNAs at baseline were associated with weight change after 2 years.

**RESULTS:** In fully adjusted models that included relevant covariates, seven miRs (i.e., miR-126, miR-15a, miR-192, miR-23a, and miR-27a) were statistically associated with weight over 2 years. MiR-197 and miR-320a remained significant after adjustment for multiple comparisons. Baseline levels of let-7f, miR-17, and miR-320c were significantly associated with 3% weight loss after 2 years in fully adjusted models.

**DISCUSSION:** This study provided evidence for longitudinal relationships between circulating microRNAs and weight. Because microRNAs characterize the combined effects of genetic determinants and responses to behavioral determinants, they may provide insights about the etiology of overweight and obesity in the context or risk for common, complex diseases. Additional studies are needed to validate the potential genes and biological pathways that might be targeted by these microRNA biomarkers and have mechanistic implications for weight loss and disease prevention.

## INTRODUCTION

In the United States, 43% of the adult population is obese.(1) Overweight and obesity are associated with substantial increased risk for at least eight of the 10 leading causes of death in the US, including type 2 diabetes (T2D).(2) Given the adverse health consequences associated with overweight and obesity, achieving and maintaining a healthy weight is a primary approach to health promotion and T2D prevention.(3) While the health risks associated with overweight and obesity have been well described for decades, national initiatives to achieve healthy weight in all individuals have not been successful, and there are vast disparities in the prevalence of overweight and obesity in the United States by race and ethnic group, geographic region, socioeconomic status, and other demographic characteristics.(1) In fact, while body mass index (BMI) has long been the standard measure of overweight and obesity, recent evidence and consensus suggest that health disparities may be perpetuated by the use of BMI as the sole measure of body composition.(4) However, while optimal measures of body composition for use in clinical practice are still being evaluated, recommendations on weight loss for disease prevention in individuals who carry excess adipose tissue are still applicable.

One of the major challenges to weight loss and maintaining a healthy weight is the complex etiology that includes contributing factors ranging from individual-level characteristics like genetic determinants to societal-level factors like access to healthy foods and safe outdoor spaces.(5, 6) In addition, there are inter-generational effects on weight in which an offspring’s weight can be determined in part by exposures *in utero*.(7) Given the limited ability to modify some of the domains of risk at extreme ends of the risk spectrum (i.e., individual genetic risk on one end and social determinants on the other end), there is a focus on interventions that target more modifiable risk factors like lifestyle, and in particular, diet and physical activity. Therefore, these lifestyle changes are the first recommended approach for weight loss for a majority of overweight and obese individuals. Structured intensive lifestyle interventions, like the one tested in the landmark Diabetes Prevention Program (DPP) trial, are effective for weight loss and showed a 58% lower incidence of type 2 diabetes (T2D) after 2 years compared to a placebo treatment.(8) However, given that these structured intensive interventions only impact a portion of the spectrum of risk domains, not all individuals achieve their weight loss goals, and among those who do, not everyone is able to sustain their weight loss. For example, after 10 years of follow-up in the Diabetes Prevention Program (DPP) trial, there was no difference in weight change between the three trial arms.(9)

One tactic for overcoming the barriers to intervening across the spectrum of risk factors related to weight loss is by improved ability to characterize the underlying physiological mechanisms related to changes in weight. Biomarkers associated with changes in weight over time may inform these mechanisms, regardless of whether they are stimulated by individual-, environmental-, or social-level factors. MicroRNAs (miRs) are short (i.e., 18-26 nucleotide) regulatory elements of the process of translation of messenger RNA (mRNA) to amino acids. Because miRs regulate gene expression, they operate as a function of both underlying genetic risk for disease as well as environmental and social factors. Circulating miRs found in serum and plasma are easily collected from blood by a standard venipuncture and are potential biomarkers associated with the mechanisms that underlie changes in weight. A number of prior studies described circulating miRs related to overweight and obesity.(10) However, a majority of studies have been cross-sectional and therefore cannot differentiate between miRs that may be associated with likelihood of future weight loss, mechanisms underlying successful weight loss, and consequent health benefits of weight loss. In addition, few prior studies were imbedded with randomized clinical trials that tested interventions to target weight loss. We have previously shown that circulating miRs show longitudinal changes in relation to fasting blood glucose in overweight participants at risk for T2D, and that these associations are affected by lifestyle interventions.(11) In addition, our prior studies have shown that these miRs target genes located in biological pathways that are known to cause T2D.(12, 13) The purpose of this study was to evaluate the longitudinal relationships between miRs and weight in individuals at risk for T2D who participated in the DPP trial. In addition, we compared miRs associated with longitudinal measures of weight to miRs measured at baseline that were associated with a weight loss outcome after 2 years to differentiate potential prognostic from mechanistic biomarkers.

## RESEARCH DESIGN AND METHODS

### Participants and Study Design

This study was a secondary analysis of data and biospecimens from participants in the DPP trial that tested two approaches to prevention of T2D. DPP sample characteristics, trial design, and methods have been described in detail previously.(14-16) Participants were recruited from 27 centers across the US between 1996-1999 with oversampling from racial minority groups.(16) Inclusion criteria included age >25 years, BMI >24 kg/m^2^, fasting glucose between 95 and 125 mg/dL, and 2-hour post-challenge glucose between 140 and 199 mg/dL. Exclusion criteria included use of medications known to alter glucose tolerance or serious illness. A total of 3,234 participants were enrolled. The study described in this manuscript includes a randomly selected subset (n=150) equally stratified by intervention arm. We included data and biospecimens collected at basline, 1-year, and 2-years in the DPP trial to evaluate longitudinal relationships between miRs and weight.

### Demographic and Clinical Data Collection

In the DPP trial, demographic characteristics, medical history, and fasting blood glucose were collected by trained study personnel at the first screening visit.(16) Physical measurements, oral glucose tolerance tests, behavioral data, and other laboratory tests were collected at the second interview screening visit.(16) History and physical exams were collected at the third screening visit.(16) Blood samples were collected and stored for future analyses. Blood was collected into vacutainers containing the preservative heparin by venipuncture. The study described in this manuscript used these existing demographic and clinical data from the DPP trial and banked biospecimens from the National Institute of Diabetes, Digestive Disease, and Kidney Diseases biorepository.

### Molecular Data Collection

The Fireplex Multiplex Circulating MicroRNA Assay (Abcam, MA) was used for direct quantification of 58 miRs from plasma. MiRs were measured from three timepoints in the DPP trial: baseline, year-1, and year-2. MiRs were hybridized to complementary oligonucleotides covalently attached to encoded hydrogel microparticles. The bound target was ligated to oligonucleotide adapter sequences that serve as universal PCR priming sites. The miR-adapter hybrid models were then denatured from the particles and reverse transcription polymerase chain reaction (RT-PCR) was performed using a fluorescent forward primer. Once amplified, the fluorescent target was rehybridized to the original capture particles and scanned on an EMD Millipore Guava 6HT flow cytometer (Merck KGaA Darmstadt, Germany).

### Statistical Analyses

The subset of participants in this study were randomly selected from the DPP cohort, equally stratified by intervention arm. All statistical analyses were performed using R (version 4.3.1).(17) Data were summarized using descriptive statistics (means and standard deviations for continuous variables and counts and percentages for categorical variables). To compare between trial arms, we used Chi-squared (categorical variables) or analysis of variance (ANOVA) (continuous variables) to evaluate the demographic and clinical characteristics of the participants. Box and whisker plots were created to show distributions of weight loss after 2 years overall and by trial arm.

Expression of individual miRs was normalized using the set of miR probes (i.e., miR-92a-3p, mir-93-5p, miR-17-5p) identified by the geNorm algorithm for each experiment.(18) A lower limit of quantification (LLOQ) equal to two time the minimum detectable difference for each miR was applied to the samples, and sub-LLOQ values were filtered. MiRs detected in ≥90% of samples were retained for comparison, then converted into z-scores to account for batch effects between miR data collection timepoints. This resulted in 35 miRs retained for analysis.

Linear mixed models were created to evaluate the relationships between individual miRs and measures of weight at three timepoints (i.e., baseline, year-1, year-2). We included a uniform set of covariates in all fully adjusted models: age, gender, race and ethnicity, baseline weight, and trial arm. Models were fit using the lme4 package for R.(19) Significance was calculated using the lmerTest package for R which applies Satterthwaite’s method to estimate degrees of freedom and generate p-values for mixed models.(20) One purpose of this study was to compare which miRs show longitudinal associations with weight to miRs measured at baseline that are associated with weight loss after 2 years. Therefore, logistic regression was used to determine which miRs at baseline predicted a 3% weight loss at the 2-year timepoint. The same uniform set of covariates was included in fully adjusted logistic regression models. Correction for multiple testing was done by applying the false discovery rate. Statistical significance was determined by a p-value<0.05.

We determined the messenger RNA (mRNA) targets and associated biological pathways of the miRs that were found to have significant associations with weight in linear mixed models. Based on our prior review of tools for target prediction(21), miRTarBase (version 9.0) was used.(22) The *Homo sapiens* microRNA-target interaction (MTI) database was downloaded from miRTarBase. We used R(17) to filter for the mRNA gene targets of the identified miRs. Filters for the search included strong experimentally validated (i.e., Western Blot, qRT-PCR, and reporter assay) evidence. Within the OrganismDbi v1.36.0, we used the Homo.sapiens package (v1.3.1) to determine the entrez IDs for target mRNAs, which were added to the results from miRTarBase. Pathway enrichment analysis was then employed to assess for overrepresentation of the mRNA targets of miRs using the Kyoto Encyclopedia of Genes and Genomes (KEGG)(23) annotation database and clusterprofiler package v4.2.2.

## RESULTS

In this random subset of 150 participants from the parent DPP trial, there were no statistically significant differences between the trial arms at baseline (**Table 1**). There were also no differences between the participants in this subset and the full DPP sample (data not shown). Overall, a majority of the sample were female (73%), White (63%), had mean age of 51 ± 10 years, and were overweight or obese (BMI 33.7 ± 6.9 kg/m^2^). Additional demographic and clinical characteristics are shown in **Table 1**. The mean weight loss in the sample overall at the 2-year timepoint was 2.8 kg (standard deviation (SD) 6.7 kg) (**Figure 1**). In the intensive lifestyle arm, the mean weight loss was 6.4 kg (SD 7.2 kg); in the metformin arm the mean weight loss was 2 kg (SD 5.2 kg) and in the placebo group, mean weight gain was 0.2 kg (SD 6.0 kg) (**Figure 1**). Weight loss differed significanty between trial arms (p < 0.001).

**Table 1.** Demographic and Clinical Characteristics.

|  | **Overall**<br>**(n=150)** | **Lifestyle**<br>**(n=50)** | **Metformin**<br>**(n=50)** | **Placebo**<br>**(n=50)** | **p-value** |
| --- | --- | --- | --- | --- | --- |
| **Age (years)** | 51 ±10 | 52 ±11 | 51 ±11 | 50 ±11 | 0.485 |
| **Gender n (%)** |  |  |  |  | 0.791 |
| **Female** | 109 (73) | 36 (72) | 35 (70) | 38 (76) |  |
| **Male** | 41 (27) | 14 (28) | 15 (30) | 12 (24) |  |
| **Race and Ethnicity n (%)** |  |  |  |  | 0.898 |
| **White** | 94 (63) | 29 (58) | 33 (66) | 32 (64) |  |
| **Black** | 28 (19) | 11 (22) | 9 (18) | 8 (16) |  |
| **Hispanic** | 22 (15) | 9 (18) | 6 (12) | 7 (14) |  |
| **Other** | 6 (4) | 1 (2) | 2 (4) | 3 (6) |  |
| **Body mass index (kg/m<sup>2</sup>)** | 33.3 ±6.6 | 32.5 ±6.3 | 33.1 ±6.6 | 34.2 ±6.8 | 0.412 |
| **Fasting glucose (mg/dL)** | 104 ±7 | 105 ±7 | 106 ±7 | 106 ±7 | 0.769 |
| **Fasting insulin (uU/mL)** | 25 ±14 | 22 ±13 | 27 ±16 | 25 ±12 | 0.152 |
| **Triglycerides (mg/dL)** | 165 ±107 | 148 ±60 | 173 ±131 | 172 ±117 | 0.426 |
| **Total cholesterol (mg/dL)** | 205 ±38 | 207 ±42 | 201 ±38 | 208 ±36 | 0.627 |
| **HDL-cholesterol (mg/dL)** | 47 ±13 | 50 ±15 | 46 ±10 | 47 ±14 | 0.372 |
| **LDL-cholesterol (mg/dL)** | 125 ±32 | 127 ±34 | 121 ±33 | 128 ±32 | 0.516 |
| **Waist circumference (cm)** | 102 ±14 | 101 ±14 | 103 ±17 | 103 ±13 | 0.850 |
| **Hip circumference (cm)** | 113 ±17 | 111 ±11 | 112 ±11 | 116 ±24 | 0.293 |
| **Average systolic blood pressure (mmHg)** | 125 ±16 | 126 ±16 | 125 ±15 | 124 ±17 | 0.846 |
| **Average diastolic blood pressure (mmHg)** | 78 ±9 | 78 ±9 | 77 ±9 | 78 ±9 | 0.869 |

**Figure 1.**
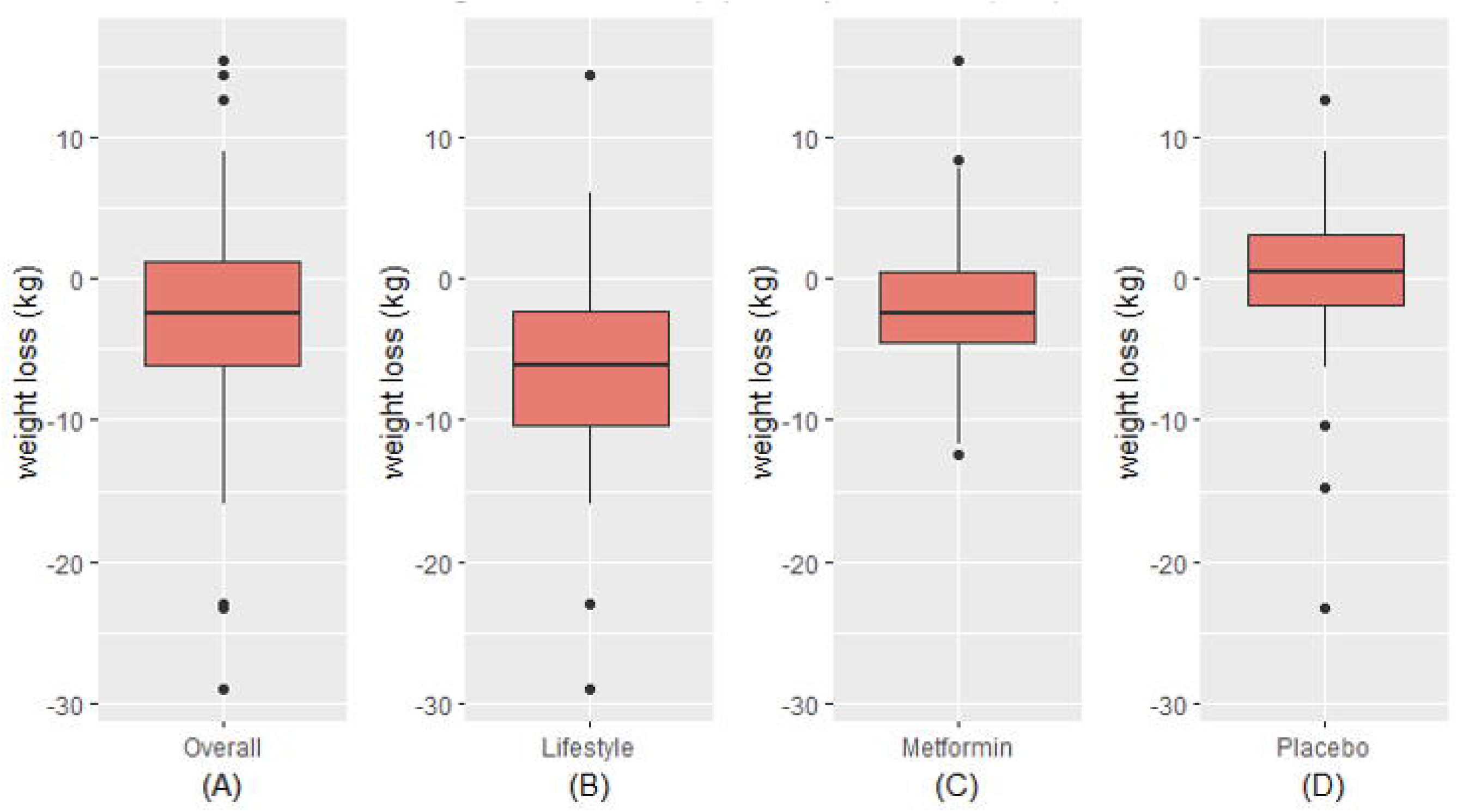
Distribution of Weight Loss Overall (A) and By Trial Arm (B-D) Within each red box, the bottom border represents the 25th percentile, the center line represents the 50th percentile, and the upper border represents the 75th percentile. The lowest point on the horizontal line represents the minimum value, and the highest point on the horizontal line represents the maximum value. Small black circles represent outliers.

Of the 35 miRs included in the analysis, seven (i.e., miR-126, miR-15a, miR-192, miR-197, miR-23a, miR-27a, miR-320a) were significantly associated with measures of weight over time (**Table 2**), though these associations did not remain significant after correction for multiple comparisons. After adjusting for relevant covariates (age, gender, race and ethnicity, baseline weight, trial arm), all seven remained statistically associated with weight, two of which (miR-197, miR-320a) remained significant in the fully adjusted models after correction for multiple comparisons.

**Table 2.** Linear Mixed Models for MicroRNAs and Weight.

| microRNA | Unadjusted models |  |  |  | Fully Adjusted models |  |  |  |
| --- | --- | --- | --- | --- | --- | --- | --- | --- |
|  | Coefficient | 95% CI | p-value | p-value adjusted* | Coefficient | 95% CI | p-value | p-value adjusted* |
| miR-126 | 0.55 | 0.03, 1.08 | 0.037 | 0.186 | 0.32 | -0.12, 0.75 | 0.147 | 0.571 |
| miR-15a | -0.77 | -1.38, -0.16 | 0.013 | 0.098 | -0.54 | -1.04, -0.02 | 0.041 | 0.204 |
| miR-192 | 0.68 | 0.12, 1.23 | 0.016 | 0.098 | 0.6 | 0.15, 1.05 | 0.010 | 0.087 |
| miR-197 | 0.76 | 0.28, 1.25 | 0.002 | 0.070 | 0.67 | 0.93, 1.0 | 0.002 | **0.026** |
| miR-23a | -0.73 | -1.26, -0.20 | 0.007 | 0.098 | -0.54 | -0.99, -0.07 | 0.022 | 0.156 |
| miR-27a | -1.28 | -1.23, -0.12 | 0.017 | 0.098 | -0.47 | -0.94, 1.02 | 0.057 | 0.248 |
| miR-320a | 0.67 | 0.16, 1.18 | 0.010 | 0.098 | 0.71 | 0.27, 1.16 | 0.002 | **0.026** |

MicroRNAs and weight were measured at baseline, year 1, and year 2 of the Diabetes Prevention Program trial. Fully adjusted models included baseline measures of age, race and ethnicity, weight, and trial arm.
CI – confidence interval; miR – microRNA
\*p-values were adjusted for multiple comparisons using the False Discovery Rate

We also tested for associations between microRNAs measured at baseline and achieving 3% weight loss after 2 years (**Table 3**). Four miRs (i.e., let-7f, miR-17, miR-30a, miR-320c) showed significant associations with 3% weight loss in unadjusted models, and all except miR-30a were also significantly associated with 3% weight loss in fully adjusted models. Neither the unadjusted nor adjusted models remained significant after correction for multiple comparisons. None of these microRNAs overlapped with those that were significant in longitudinal models, though miR-320a and miR-320c are isoforms.

**Table 3.** Odds of Achieving 3% Weight Loss After Two Years.

| Unadjusted Models |  |  |  |  | Fully Adjusted Models |  |  |  |
| --- | --- | --- | --- | --- | --- | --- | --- | --- |
| miR | Odds Ratio | 95% CI | p-value | *p-value<br>adjusted | Odds Ratio | 95% CI | p-value | *p-value<br>adjusted |
| let-7f | 1.96 | 1.19, 3.34 | 0.010 | 0.321 | 2.04 | 1.13, 3.85 | 0.022 | 0.253 |
| miR-17 | 1.41 | 1.02, 2.00 | 0.037 | 0.321 | 1.40 | 0.94, 2.14 | 0.105 | 0.360 |
| miR-30a | 1.59 | 1.07, 2.46 | 0.027 | 0.321 | 1.48 | 0.96, 2.39 | 0.088 | 0.360 |
| miR-320c | 1.47 | 1.05, 2.11 | 0.029 | 0.321 | 1.86 | 1.23, 3.02 | 0.006 | 0.223 |

MicroRNAs were measured at the baseline timepoint in the Diabetes Prevention Program trial. Fully adjusted models included baseline measures of age, gender, race and ethnicity, weight, and trial arm.
CI – confidence interval; miR – microRNA
\*p-values were adjusted for multiple comparisons using the False Discovery Rate method

We identified four mRNAs that have strong functional evidence from the miRTarBase database(22) to be targeted by the miRs that showed longitudinal associations with weight: BCL2 apoptosis regulator (*BCL2*), CMI1 proto-oncogene polycomb ring finger (*BMI1*), forkhead box O3 (*FOXO3*), and phosphatase and tensin (*PTEN*). There were seven KEGG pathways that contain genes that are overrepresented as mRNA targets of six of the miRs identified in this study to have longitudinal associations with weight (miR-126, miR-15a, miR-192, miR-23a, miR-27a, miR-320a) (**Supplementary Table 1, Supplementary Figure 1**).(24) There were 37 KEGG pathways with overrepresented gene targets for a subset of four of the miRs that had longitudinal assocations with weight (miR-126, miR-23a, miR-27a, and miR-320a) (**Supplementary Table 1, Supplementary Figure 1**).(24) No pathways contained genes that were overrepresented targets of all seven miRs. There were three additional pathways (Platinum drug resistance, Colorectal cancer, p53 signaling pathway) that contained overrepresented genes for subsets of six of the seven miRs (**Supplementary Table 1, Supplementary Figure 1**).(24)

## DISCUSSION

Leveraging data from the DPP clinical trial, we identified the subset of miRs that show significant longitudinal associations with weight that may be potential candidates for understanding the mechanisms underlying changes in weight over time in response to the risk reduction interventions tested in DPP. We also identified which miRs measured at baseline predicted weight loss after 2 years that might be prodromal biomarkers for weight loss, as we have previously shown for incident T2D.(25) With the exception of two miR isoforms of miR-320, there was no overlap between the miRs that showed longitudinal versus predictive associations with weight. This observation validates the supposition that some miRs may be useful clinical biomarkers to provide both earlier and more individualized risk prediction while other miRs may be potential therapeutic targets to prevent development of T2D and related complications.

Most of the prior research on miRs in humans has been either cross-sectional or longitudinal measures were done only for disease-related outcomes but not for miR biomarkers. Because miRs regulate expression of mRNAs in order to control biological processes, they may be implicated in risk for and progression to disease, the manifestation of disease, and disease-related complications. Prior studies that did not measure miRs serially cannot differentiate where in this trajectory of disease development and sequelae a given miR may be actively regulating mRNA expression. For example, one of the first studies focused on miR biomarkers related to T2D identified differential expression of miR-126 in individuals who developed T2D after 10 years.(26) This study provided evidence that miR-126 may be a useful prodromal biomarker to identify who is at highest risk for incident diabetes. However, follow-up studies showed that miR-126 is related to endothelial cell function, and therefore this miR may be related to complications from, rather than development of T2D. To further explore the potential mechanistic roles of miRs, *in vitro* studies have provided evidence for the functional relationships between miRs and their mRNA targets and resulting gene products, however this highly controlled environment may not fully characterize the activity of a given miR in a complex human environment.

In the present study, miR-197 showed significant longitudinal associations with weight. Our own prior studies also identified miR-197 in an assessment of longitudinal expression levels of miRs with fasting blood glucose measured after 1 year in overweight and obese participants from clinical trial that assessed the impact of a behavioral intervention on T2D risk factors.(11) A second study also identified miR-197 as a member of a factor of miRs that target genes and pathways related to inflammation and metabolism in individuals at risk for T2D.(12) Excess adipose tissue is widely known to be related to increased systemic inflammation.(27) A study focused on osteoarthritis described functional relationships in chondrocytes *in vitro* between miR-197 and inflammatory markers (i.e., interleukin (IL)-β, IL-6, and tumor necrosis factor (TNF)-α).(28) Another study reported that miR-197 was present in biopsied human omental and subcutaneous adipose tissue, consistent with a shared embryonic histologic origin, but the levels were significantly higher in the omental tissue compared to subcutaneous (fold change (FC) 1.2, p=0.04)),(29) suggesting differing biologic activity between the two sub-types. In support of the evidence that miR-197 may regulate adipocyte differentiation, a prior study that assessed the impact of miR-197 delivered to breast cancer cells *in vitro* found inhibition of expression of peroxisome proliferator-γ (*PPARγ*) mRNA and protein.(30) *PPARγ* is highly expressed in adipocytes and regulates differentiation of preadipocytes.(31) In overrepresentation analysis in the current study, common mRNA targets of miR-197 included *IL-8, FOXO3*, and mitogen activated protein kinase 1 (*MAPK1*), which map to pathways related to cancer, inflammation/immunity, and cellular activity (**Supplementary Table 2**).(24) While relatively little is known about the function of miR-197 in relation to body weight in human tissues *in vivo*, prior evidence from other phenotypes suggests that this miR may be involved in processes that underlie regulation of body weight.

Among the miRs that showed significant longitudinal relationships with weight, miR-15a, miR-23a, and miR-27a were included in the same previously described factor of miRs that was derived by assessing statistical associations between miRs measured at five timepoints over 1 year.(12) These measures were taken from overweight and obese participants at risk for T2D who participated in a behavioral intervention trial.(32) Overrepresentation analysis in the prior and the current studies showed that the miRs in this factor target mRNAs and pathways related to inflammation and metabolism, two known mechanisms that underlie risk for T2D (**Supplementary Table 1**).(24) In contrast, in our prior study of participants from the DPP that investigated miRs for prediction of incident T2D, none of the miRs with longitudinal relationships were identified as optimal predictive biomarkers. This is further support for the notion that miRs may have discrete roles as biomarkers for risk prediction as opposed to specific mechanisms that underlie development of T2D and related complications.

While we hypothesize that miRs that are predictive biomarkers are not completely intersecting with miRs biomarkers that characterize underlying mechanisms, we also suggest that for some miRs, there may be overlap in the information that can be inferred and potential clinical utility. Our prior study of participants from the DPP cohort identified *BCL2* as a mRNA target of miRs that were optimal predictors of incident T2D.(13) The current study also identified *BCL2* as a predicted target of miR-126, miR-15, miR-192, and miR-23a that have longitudinal associations with weight (**Supplementary Table 2**).(24) Another example is miR-320. The current study identified longitudinal relationships between miR-320a and weight as well as a predictive relationship between its isoform, miR-320c, and 3% weight loss after 2 years. MiR isoforms, or isomiRs are nearly alike molecules that typically have identical seed sequences but can have small variations in length or nucleotide sequence that allow for highly specified regulatory activity.(33) A handful of prior studies reported relationships between miR-320 and obesity and T2D related conditions. MiR-320a has been shown to increase in response to a high fat diet in mice, whereas miR-320c was unchanged.(34) When miR-320a mimics were administered to pancreatic INS1 β cells *in vitro*, decreased proliferation and increased apoptosis and reactive oxygen species were observed, suggesting this isoform may be damaging for pancreatic β cells and a potential therapeutic target for T2D prevention.(34) Another study showed that miR-320 regulated hepatic adiponectin signaling in rats who underwent duodenal-jejunal bypass surgery for weight loss, though the specific isoform was not reported.(35) Two other studies reported that miR-320 regulates endothelial function, endoplasmic reticulum stress, and inflammation *in vitro*, though again, the specific isoforms studied was not reported.(36, 37) Future longitudinal studies in humans, followed by functional validation studies, are needed to serially measure miR-320 isoforms to determine if this miR is a prodromal predictor of T2D, related to mechanisms underlying risk for and development of T2D, or both, and whether this differs by isoform.

MiR-192 is the fifth miR that showed significant associations with weight in fully adjusted models. Our prior study of overweight and obese individuals at risk for T2D identified significant longitudinal associations between this miR and fasting blood glucose after 1 year.(11) MiR-192 measured in visceral adipose tissue from obese individuals has previously been shown to have associations with triglycerides (r=-0.387; p=0.046), high-density lipoprotein (HDL) cholesterol (r=0.396; p=0.041), and BMI (r=0.396; p=0.041).(38) Subsequent *in vitro* experiments showed that this miR targets genes known to regulate adipocyte differentiation and lipid homeostasis.(38) Another study focused on renal ischemia/reperfusion injury showed that miR-192 regulates expression of the fat mass and obesity (*FTO*) mRNA.(39) MiR-192 showed evidence for overrepresented mRNA targets in three unique KEGG pathways in the present study: ECM-receptor interaction, Hypertrophic cardiomyopathy, and Dilated cardiomyopathy (**Supplementary Table 2**).(24) We previously reported that a factor of co-expressed miRs related to fasting blood glucose in overweight and obese individuals also showed overrepresentation of mRNA targets in the latter two pathways.(12) MiRs associated with 3% weight loss after 2 years also have some evidence for associations with risk for T2D or related complications. Let-7f has largely been reported in studies on vascular and autoimmune conditions. This miR was included in a factor of miRs in our prior study of overweight and obese people at risk for T2D, and collectively, an overrepresentation analysis showed that the miRs in this factor target mRNAs and pathways related to endocrine and hormone function.(12) MiR-17 is most commonly studied in relation to cancer, which has elucidated evidence for targeting of *PTEN* in cancer cells(40), and the PTEN protein has well-described associations with obesity and insulin sensitivity(41) and is a predicted target of miR-23a, miR-27a, and miR-320a, which showed longitudinal associations with weight in the study described here. MiR-17, which predicted 3% weight loss, was previously associated with change in fasting blood glucose over 1 year in overweight or obese people at risk for T2D.(12) Finally miR-30 has primarily been studied in relation to cancer as well, however several of these studies reported regulation of various inflammatory pathways in cancer cells.

### Limitations

Given the number of miR biomarkers tested, we selected a standard set of covariates to be included in all models. There may be additional demographic or clinical characteristics that covary in relation to a specific miR, including clinical characteristics that are on the causal pathway for changes in weight, which were not captured in this analysis. Many miRs have been described and studied in relation to a specific disease condition (e.g., a single type of cancer). While this study was focused on weight in individuals at risk for T2D, overweight and obesity are also major risk factors for cancer and many other common, complex diseases. The findings in this study sample may also have relevance for people with other diseases that are more likely in the setting of overweight and obesity.

We described miRs that change over time in relation to weight in participants from the DPP trial. This is the largest study, to our knowledge, the relate these changes over time as opposed to cross-sectional or predictive longitudinal study designs. Some identified miRs have known associations with mechanisms related to overweight and obesity (e.g., inflammation, adipocyte differentiation) while the mRNA and biological pathway targets of other miRs are not well understood. We also showed that miRs that may be predictive biomarkers are not necessarily the same miRs that correspond with the underlying mechanisms of weight at multiple time points. Future studies that validate these mechanistic relationships may provide new potential therapeutic targets for weight loss and prevention of T2D and other common, complex diseases.

## Figure Legends

**Supplementary Figure 1**. This upset plot depicts the overlap in the pathways that were enriched for mRNA targets of the 7 microRNAs that were significantly associated with weight over 2 years. Intersection size is the number of pathways targeted by the combination of microRNAs indicated by black dots below. Numbers above dark grey bars represent the intersection size value. Set size is the total number of pathways targeted by each individual microRNA.

## DATA AVAILABILITY

Some or all datasets generated during and/or analyzed during the current study are not publicly available but are available from the corresponding author on reasonable request.

