## Supplementary figures and images for "Longitudinal Associations Between MicroRNAs and Weight in the Diabetes Prevention Program"

### Table 1.DOCX

**Supplementary Figure 1.** Upset Plot for Predicted KEGG Pathways by Combination of MicroRNAs

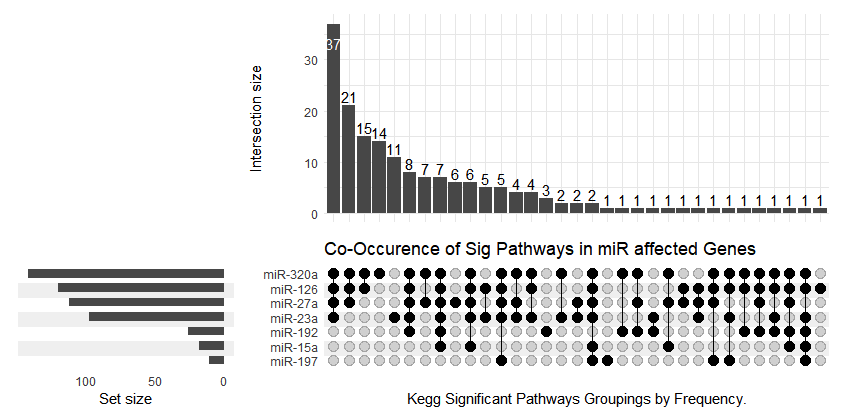
